# Sexual selection and pseudogenization in primate fertilization

**DOI:** 10.1101/2022.05.16.491899

**Authors:** Jolie A. Carlisle, Andrew L. Bickell, Michael W. Hart

**Affiliations:** Cornell University, Department of Molecular Biology and Genetics; Simon Fraser University, Department of Biological Sciences

**Keywords:** Neofunctionalization, sexual conflict, mating system, speciation, infertility

## Abstract

The mouse sperm protein ZP3R interacts with proteins in the egg coat and mediates sperm–egg adhesion in a species-specific manner. Understanding the function and evolution of such genes has important implications for studies of speciation, reproductive success, and infertility. A recent analysis showed that (1) the human pseudogene *C4BPAP1* is the ortholog of *Zp3r*, (2) *ZP3R* pseudogenization evolved independently in parallel among several primate lineages, and (3) functional *ZP3R* genes evolve under positive selection among other primate species. The causes of this pseudogenization of *ZP3R* are unknown. To explore one plausible cause (changes in sexual selection on males), we searched for *ZP3R* pseudogenes in recently published genomes, then compared sexually selected male traits among lineages with and without a functional *ZP3R*. We found that traits associated with more intense sexual selection on males (large male body size, larger male canines, larger testes) were associated with functional *ZP3R* expression, and suggest that a relaxation of sexual selection may be associated with selection for *ZP3R* pseudogenization. This proposed causal relationship implies an evolutionary cost to maintaining redundancy in the suite of primate fertilization genes.

**Lay summary:** In sexual interactions more is often assumed to be better. But the evolution of animal genomes suggests that sometimes less is more: the adaptive loss of genes that function in sex may be favored by selection. How could this happen? One surprising answer comes from analyzing humans and some other primate species that have turned off a key gene called *ZP3R* that helps sperm bind to eggs. The loss of that gene function in some primates is associated with male traits (smaller bodies, smaller canines, smaller testes) that often indicate less vigorous selection on males to compete for matings with females. That correlation implies that the same selection acting on male morphological traits may also act on sperm molecular traits. The correlation also implies that it’s expensive to keep some genes turned on, and that when they’re no longer helpful it’s adaptive to turn them off or allow them to become fallow. This economy of gene expression in sex is an under-explored area of research.

## Introduction

Successful fertilization is critical for all sexually reproducing organisms. Interactions between sperm and egg in animals are mediated in part by a suite of proteins that are expressed on the surfaces of gametes and mediate several types of cellular interactions at the time of fertilization, ranging from sperm chemoattraction to gamete fusion (Hirohashi et al., 2008). The genes that encode these gamete surface proteins are often under strong selection (Metz & Palumbi, 1996; Vacquier, 1998; Swanson & Vacquier, 2002). Understanding how selection shapes the evolution of gamete interaction can potentially illuminate the phenotypic and genetic causes of fertility variation and reproductive compatibility between mates, and the evolution of reproductive isolation between populations and species. Understanding the molecular basis for sperm–egg interactions in mammals is of particular interest for the development of both contraceptives (Meeusen et al., 2007) and infertility treatments (Huang et al., 2014).

Although many important molecules have been described for their role in mammalian fertilization (e.g., sperm–egg plasma membrane fusion; Inoue et al., 2013; Allingham & Floriano, 2021; Brukman et al., 2022), the catalogue of genes and proteins that mediate fertilization can be incomplete (Carlisle & Swanson, 2021). That gene repertoire evolves in part by whole-gene duplication, the addition of new paralogs into related fertilization functions, functional redundancy of some genes (Ikawa et al., 2010; Avella et al., 2013; Lu et al., 2019), and the loss of functional members of such gene families (recently reviewed by Carlisle et al., 2022; Rivera & Swanson, 2022). For example, the mouse egg coat proteins ZP2 and ZP3 interact with mouse *Zp3r* (or *Zona pellucida 3 receptor*, also known as *Sp56*) expressed in the sperm acrosome (Buffone et al., 2008; Wassarman, 2009). However, male *Zp3r*-/ *Zp3r*-null homozygotes are fertile (Muro et al., 2012), probably because *Zp3r* is functionally redundant with other male-expressed genes and proteins that bind eggs. Functional redundancies in fertilization may evolve as part of the arms race between males and females driven by sex differences in optimal values for reproductive traits including the specificity or strength of sperm–egg binding interactions (Gavrilets, 2000; Turner et al., 2008).

The human ortholog of mouse *Zp3* has long been well known (Buffone et al., 2008), but until recently the identity of its human ortholog was uncertain. The RCA gene cluster on mouse chromosome 1 (chr1) includes *Zp3r* and several other members of the C4b-binding protein family of genes and pseudogenes (McLure et al., 2004) that encode regulators of complement activation in the innate immune system (Arenzana et al., 1995). Mouse *Zp3r* is descended from mouse *C4bp* by an ancient gene duplication event, and *Zp3r* has acquired a novel pattern of expression and protein localization (in the testis and acrosome) and a novel function (in sperm binding to the egg coat).

The human RCA cluster includes only one functional gene (*C4BPA*) and a non-functional pseudogene (*C4BPAP1*). Several previous population genetic studies assumed that the functional gene *C4BPA* is the human ortholog of *Zp3r* (Rohlfs et al., 2010; Hart et al., 2018; Morgan & Hart, 2019). Carlisle et al. (2024) analyzed synteny between genes in the mouse and human RCA clusters, and phylogenetic relationships among the members of the C4b binding protein gene family, to show that *C4BPAP1* is the human ortholog of mouse *Zp3r*. Expression of that human *ZP3R* pseudogene mRNA in the testis (Carithers & Moore, 2015) is consistent with its former function in fertilization.

Carlisle et al. (2024) also identified *ZP3R* pseudogenization events that occurred independently across some but not all other primate lineages. The parallel loss of *ZP3R* function was unambiguously inferred from differences in the genomic locations and types of frameshift mutations and premature stop codons among *ZP3R* coding sequences from eight other lineages. Given the evidence for positive selection acting on divergence among functional primate *ZP3R* genes (Carlisle et al., 2024), and the general tendency for sexual selection to cause rapid divergence of genes that mediate reproductive success (Gavrilets, 2000; Uy & Borgia, 2000; Van Doorn et al., 2001; Swanson & Vacquier, 2002; Patiño et al., 2016), Carlisle et al. (2024) concluded that “recurrent, lineage-specific gene loss events [are] suggestive of strong selection for gene loss [but] it is still unclear what is driving the loss of *ZP3r* in primates.”

Here we explore evidence for one possible mechanism driving the evolutionary loss of *ZP3R* expression: changes in the strength of sexual selection on males that permits or favours the loss of gene function and reduced redundancy in sperm adhesion to the egg coat. We added to the data from Carlisle et al. (2024) by finding functional *ZP3R* coding sequences or pseudogenes in recently published genomes of 17 additional primate species. We used previously published morphological data to characterize variation among primate lineages in three sexually-selected traits, and we show that smaller males with smaller canines (but not smaller testes) are associated with *ZP3R* pseudogenization events. One interpretation of that pattern across primate lineages is that phenotypic markers of the intensity of precopulatory sexual selection predict the conditions for loss (and perhaps for gain) of gene function in fertilization.

## Methods

We used data for *ZP3R* pseudogenization from Carlisle et al. (2024), including their discoveries of functional or non-functional *ZP3R* gene structure in 30 non-human primate species (Figure 1). We used the same methods (Appendix S1) to find functional *ZP3R* genes or pseudogenes in the genomes of 17 additional species from Shao et al. (2023).

**Figure 1.**
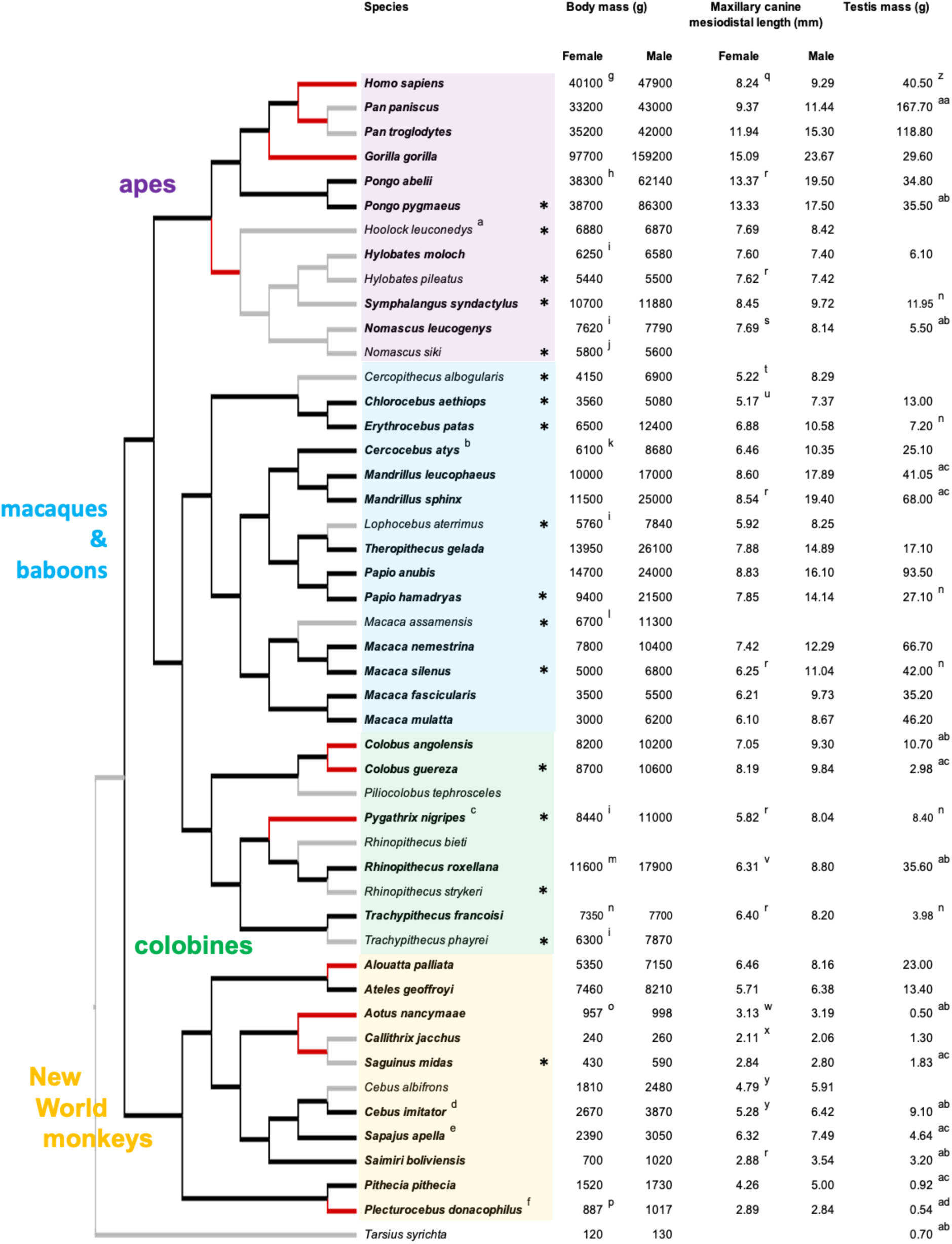
*ZP3R* functionality and morphological variables from 31 primate genomes analyzed in Carlisle et al. (2024) and 17 other primate genomes analyzed in this study (marked by asterisks). Cladogram from 10kTrees (Arnold et al., 2010). New *ZP3R* data come from genomes in Shao et al. (2023). Red edges in the cladogram show 11 parallel origins of *ZP3R* pseudogenes in humans, *Pan*, *Gorilla gorilla*, gibbons, *Colobus angolensis* and *C. guereza*, *Pygathrix nigripes*, *Alouatta palliata*, *Aotus nancymaae*, the most recent common ancestor of *Callithrix jacchus* and *Saguinus midas,* and in *Plecturocebus donacophilus*. Twelve species names in plain type show species for which some morphological data were missing; those species were not included in the principal components analysis. Grey edges in the cladogram show species or lineages that were not included in the comparative analysis (calculation of relative size as residuals from lineage regressions, BayesTraits estimate of ancestral states for relative sizes, or one-sample tests) because those species were descended from the same single pseudogenization event or because data were missing. Superscript letters indicate footnotes that explain species names or give data sources for five morphological variables that were used in quantitative analyses: a, *Bunopithecus hoolock* in 10kTrees; b, *C. torquatus atys* in 10kTrees; c, *P. nemaeus* in 10kTrees (two species were recent split, probably hybridize); d, *C. capucinus* in 10kTrees; e, *Cebus apella* in 10kTrees; f, *Callicebus donacophilus* in 10kTrees; g, body masses from Lindenfors & Tullberg (1998) unless otherwise noted; h, Markham & Groves (1990) for “Sumatran” individuals; i, Thorén et al. (2006); j, https://primate.wisc.edu/primate-info-net/pin-factsheets/pin-factsheet-white-cheeked-gibbon; k, Kennard & Willner (1941); l, Gordon (2006); m, Smith & Jungers (1997); n, Lupold et al. (2019); o, Spence-Aizenberg et al. (2018); p, Gron (2007); q, canine lengths from Thorén et al. (2006) unless otherwise noted; r, Plavcan & Ruff (2008), similar to M. Skinner (unpublished data); s, as *Hylobates concolor*; t, as *C. mitis*;u, as *Cercopithecus aethiops*; v, Pan & Oxnard (2001); w, to include this lineage in the analysis we used canine sizes for *A. trivirgatus*, which have adult body sizes similar to *A. nancymaae*; x, Leutenegger & Cheverud (1982); y, Masterson (2003); z, testis masses from Harcourt et al. (1981) unless otherwise noted; aa, Dixson (2012); ab, Ports & Jensen-Seaman (2024); ac, Harcourt et al. (1995); ad, to include this lineage in the analysis we used testis sizes for *P. cupreus* from Conley et al. (2022), in which adult body sizes and life histories are similar to *P. donacophilus* **ALT TEXT**: A phylogeny for 48 primate species with data for body mass, canine length, and testis mass. Eighteen species have evolved a *ZP3R* pseudogene; others have a functional *ZP3R*.

We analyzed the coevolution of *ZP3R* with phenotypic traits that are expected to vary with (and reflect past selection arising from) the intensity of sexual selection including male-male competition for mates or for fertilizations. We compiled data from the literature on female body mass (g), male body mass, female canine size, male canine size, and testis mass (g) from 48 species (Appendix S2). We used mesiodistal length (mm) of the maxillary (upper) canine (Thorén et al., 2006). The data include 12 species of apes, 15 species of macaques and baboons (Cercopithecinae), 9 species of colobines, 11 species of New World monkeys, and 1 tarsier (Figure 1). We interpreted the results as evidence for a correlated response to sexual selection on male morphological traits and on male molecular traits.

Cassini (2020) reviewed several life history traits that are also expected to covary with the intensity of sexual selection, including mating systems that range from social groups with monogamous male-female pairs (and limited scope for competition between males or among sperm) to social groups with multimale mating by females (and greater scope for sexual selection arising from competition for fertilizations) (see Lindenfors & Tullberg, 1998; Adret et al., 2018; Spence-Aizenberg et al., 2018). Because sexual size dimorphism is especially pronounced in primates relative to other mammals (Plavcan, 2001; Tombak et al., 2024), and traits that are closely associated with male competition for matings or for fertilizations are expected to covary with the molecular evolution of genes that encode fertilization molecules (Prothmann et al., 2012; Ports & Jensen-Seaman, 2024), we focused on those five morphological variables in our analyses of *ZP3R* pseudogenization.

For 36 species we found complete morphological data for all five variables (9 apes, 12 macaques and baboons, 5 colobines and 10 New World monkeys). For those species we summarized covariation among the five morphological variables in a principal components analysis using covariances and standardized values (Appendix S3).

We used a regression method like that of Ports & Jensen-Seaman (2024) to characterize relative male size for each of those 36 species as the residual from a linear regression against female size (male body mass = 1.61 × female body mass – 958). We estimated relative male canine length as the residual from a linear regression against female canine length (male canine length = 1.58 × female canine length – 1.11). And we estimated relative testis mass as the residual from a linear regression against male size (log[testis mass] = 0.82 × log[male body mass] – 2.11).

We then used those three traits in a comparative analysis. We found independent origins of different pseudogenizing mutations in 11 lineages (red in Figure 1; see Carlisle et al., 2024), including *Alouatta palliata*, *Aotus nancymaae*, *Colobus angolensis* and *C. guereza* (with four different unshared mutations), *Gorilla gorilla*, *Homo sapiens*, *Pygathrix nigripes*, and *Plecturocebus donacophilus* (Figure 1). We also found shared pseudogenizing mutations in the marmoset *Callithrix jacchus* and the tamarin *Saguinus midas*; in the two species of *Pan*; and in all six species of gibbons. Each of those represent a single origin of a *ZP3R* pseudogene in the most recent common ancestor of each of those species groups.

We used the phyloglm function in the R package phylolm 2.6.5 (Ho & Ane, 2014) to fit the phylogenetic logistic regression of Ives & Garland (2010) for a binary response variable (a pseudogene or a functional *ZP3R*) and a continuous predictor variable (relative male mass, relative male canine length, or relative testis mass) with 1000 bootstrap replicates. We used trait data and the phylogram with branch lengths for all 36 species from 10kTrees (the same topology shown in Figure 1; Appendix S4). We used the logistic maximum likelihood estimator in phyloglm (for canine size and testis size) or the generalized estimating equations for a Poisson regression with penalized likelihood (for male size, where the MLE method failed to converge on a solution).

A functional *ZP3R* is the ancestral state shared among primates and rodents, and pseudogenization appears to be irreversible (see Carlisle et al., 2024). Thus trait values on internal edges in the phylogeny can be associated with known *ZP3R* states, including a pseudogene that evolved in the common ancestor of *Pan* species, a pseudogene that evolved in the common ancestor of *Callithrix* and *Saguinus*, a pseudogene that evolved in the common ancestor of the gibbons (shown in red in Figure 1), and a functional gene for all other internal edges (shown in black in Figure 1). This makes the phylogenetic logistic regressions problematic because they model evolution of the dependent and independent variables along all edges of the phylogeny, including edges on which no evolution of *ZP3R* is expected and evolution of the morphological traits is not informative (e.g., the 10 edges descended from the MRCA of gibbons, which all have the irreversible derived pseudogene).

To complement those results, we added a second simpler test. We used BayesTraits v4.1.1 (Barker et al., 2007) to infer the ancestral value of the trait on internal branches or edges of the phylogeny under a random walk model using the MCMC method (Appendix S5); similar ancestral state estimates are one analytical step in the phylogenetic logistic regression analysis as well. We used the same phylogram from the phylogenetic logistic regressions.

We then used a one-sample *t* test to compare the mean and variance for each trait in the sample of edges (3) and species (8) that independently evolved a pseudogene (shown in red in Figure 1) to a test value based on the phylogeny-wide expectation for that trait (the average for all 51 other internal edges and species with a functional *ZP3R*, shown in black in Figure 1).

The one-sample tests avoid modeling trait evolution along edges with irreversible pseudogenes, including the 10 internal and terminal edges among the gibbons that all descended from a single origin of a pseudogene in their common ancestor (Figure 1), the two terminal edges leading to *Pan paniscus* and *P. troglodytes*, and the two terminal edges leading to *C. jacchus* and *S. midas*. However, the one-sample tests do not account for autocorrelation among trait values along the internal edges used to calculate the phylogeny-wide expectation.

## Results

Most pseudogenes were found in species with small males, small canines, and small testes (Figure 2). Both females and males were larger in species with a pseudogene (mean 15.6 kg, 21.0 kg, respectively) than in species with a functional *ZP3R* (8.5 kg, 14.7 kg), mainly because our sample of species (12) and observations of *ZP3R* pseudogenes (10) included many apes with relatively large body sizes (Figure 1). However, when plotted against female size, males were relatively larger among species with a functional *ZP3R* (Figure 2), and this difference was significant in other analyses (see below).

**Figure 2.**
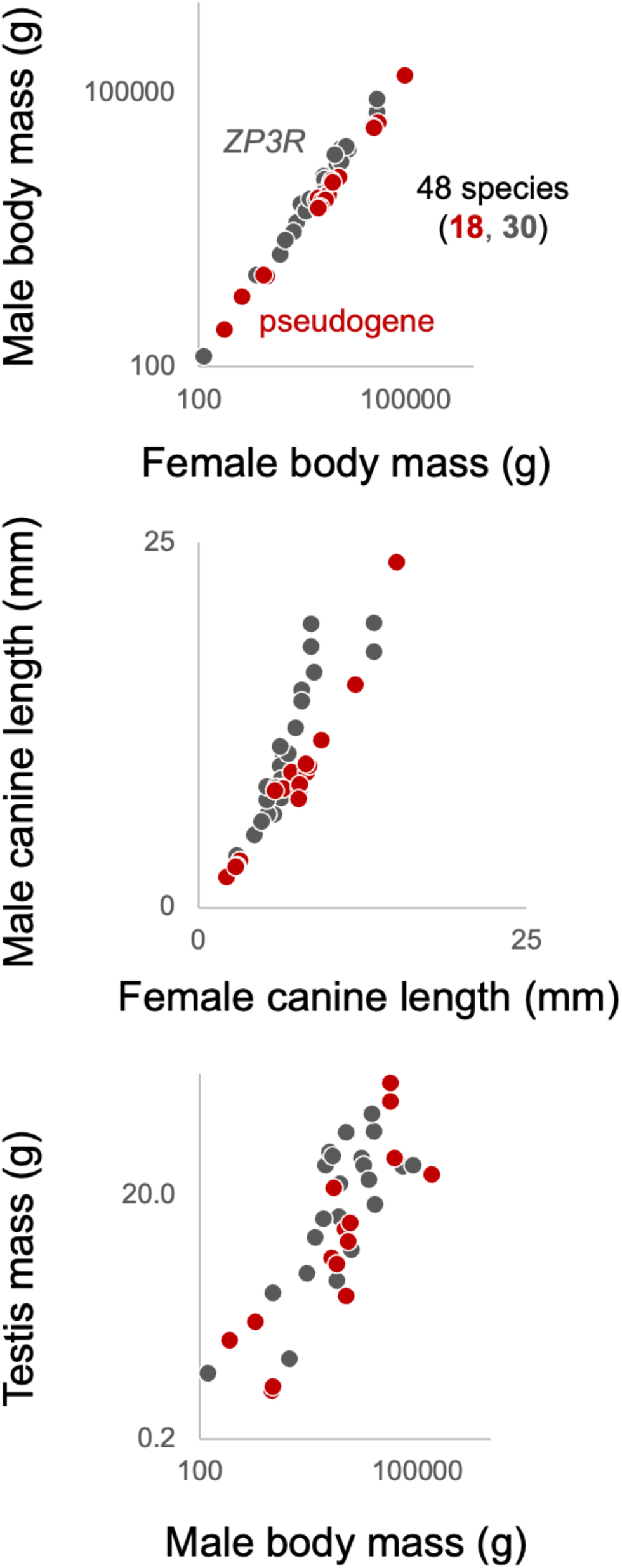
Female and male body masses (on logarithmic axes), canine lengths (on linear axes), and male body mass and testis mass (on logarithmic axes) for all 30 primate species with a functional *ZP3R* (including some species with some missing data) and all 18 species with a pseudogene (including some such as the gibbons species that share a single pseudogenization event from a common ancestor) (see Figure 1) **ALT TEXT:** Three scatter plots for sexually selected traits including male body mass plotted against female body mass, male canine length against female canine length, and testis mass against male body mass.

The pattern of smaller size associated with pseudogenes was strongest in the data for canine length (Figure 2): males of species with a pseudogene had smaller canines (mean 8.6 mm) than males of species with a functional *ZP3R* (10.7 mm); and the difference was reversed in females (7.2 vs. 6.9 mm). The difference in relative male canine length was partly caused by the large male canines found in most macaques, baboons, and other cercopithecines, in which no species was found to have evolved a pseudogene (Figure 1). Notably, that group includes many species with promiscuous or multi-male mating systems and high expected competition among males for access to females or for fertilization of eggs (Lindenfors & Tullberg, 1998; Cassini, 2020).

The association between pseudogenes and sexually selected traits was weakest in data for testis size. Males of species with (28.6 g) and without (28.4 g) a pseudogene had nearly identical mean testis mass. This similarity was caused in part by the wide variation in absolute and relative testis mass among apes that have (*Pongo*) or lack (*Homo*, *Gorilla,* gibbons) a functional *ZP3R* and in particular by the very large testes for both species of chimpanzees (Figure 1).

The principal components analysis helped to reduce the dimensionality of that variation to three simple vectors (Table 1): a compound size variable for which all five variables had positive coefficients of similar magnitude (PC1; 74% of variance); a contrast between testis mass versus male and female body mass (PC2; 16%); and a contrast between male and female canine length versus body masses and testis mass (PC3; 7%). In the space defined by PC2 and PC3, species with a functional *ZP3R* tended to have higher or positive scores for PC2 (large testes, small male and female body sizes) and PC3 (large female and male canines, small body and testis sizes) (Figure 3). As in the broader data set including all species (Figure 2), the difference was larger for PC3 (canine size relative to body size) than for PC2 (testis size) in part due to the wide variation in testis mass among apes.

**Table 1.** Principal components analysis of five morphological variables for 36 primate species. Bold values for some loadings in two eigenvectors show the variable(s) that contribute(s) most to positive scores for the two vectors that define a space in which sexually selected traits vary among species with and without a *ZP3R* pseudogene **ALT TEXT:** Results from a clustering analysis of five morphological variables from 36 primate species. Three eigenvectors explain most of the variation, including a contrast between testis size and body size, and a contrast canine size with body and testis size.

|  | PC<sub>1</sub> | PC<sub>2</sub> | PC<sub>3</sub> | PC<sub>4</sub> | PC<sub>5</sub> |
| --- | --- | --- | --- | --- | --- |
| Eigenvalue | 3.72 | 0.810 | 0.359 | 0.0868 | 0.0144 |
| % of variance | 74.5 | 16.2 | 7.19 | 1.73 | 0.288 |
| Eigenvectors |  |  |  |  |  |
| Female canine length | 0.491 | -0.0181 | **0.343** | 0.794 | -0.0902 |
| Male canine length | 0.465 | 0.0896 | **0.664** | -0.555 | 0.157 |
| Female body mass | 0.477 | **-0.273** | **-0.475** | -0.0183 | 0.685 |
| Male body mass | 0.479 | **-0.334** | **-0.327** | -0.242 | -0.701 |
| Testis mass | 0.288 | **0.897** | **-0.326** | -0.0246 | -0.0704 |

**Figure 3.**
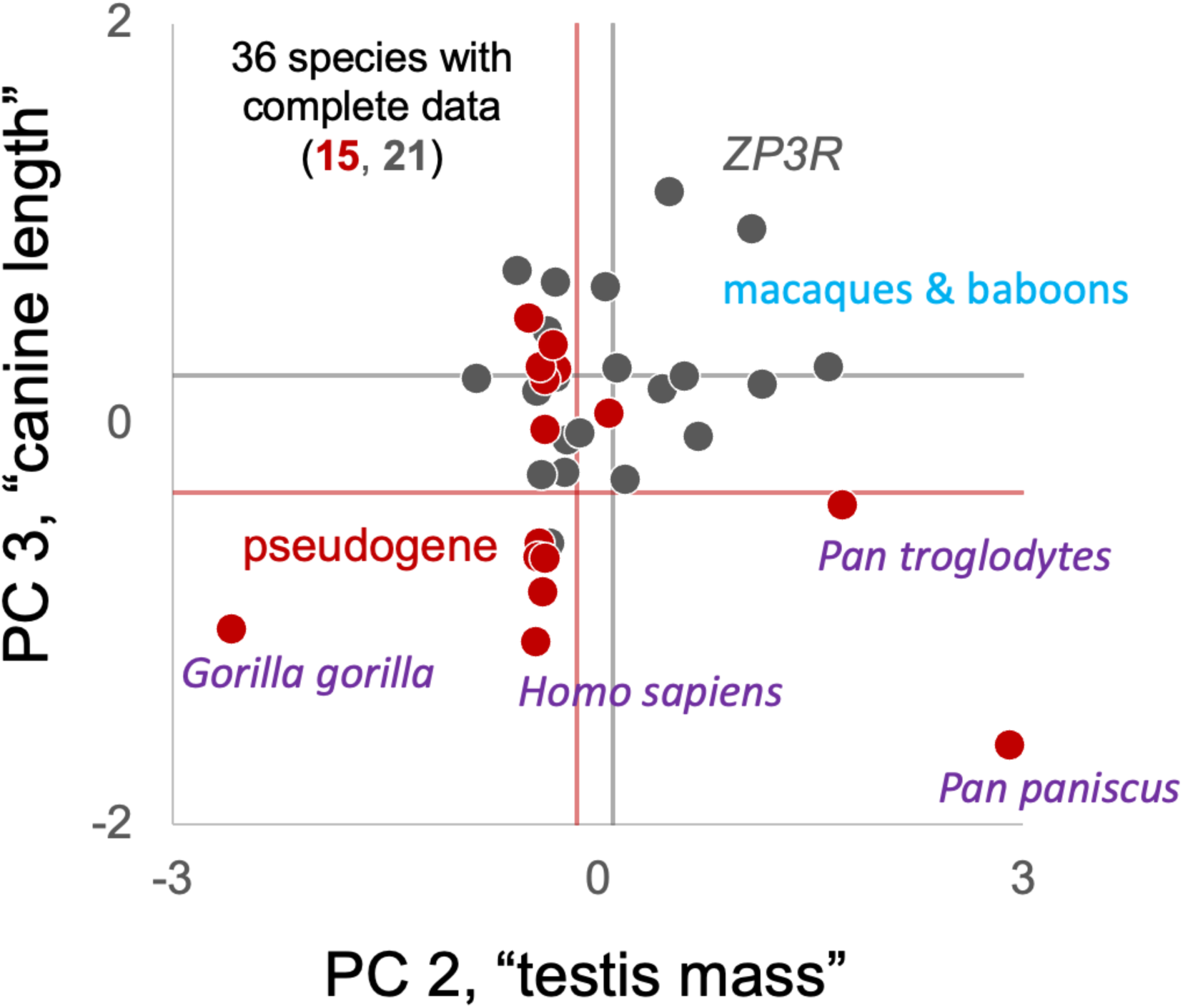
Scores on the second and third eigenvectors of a principal components analysis of five morphological variables from 21 primate species with a functional *ZP3R* (grey) and 15 species with a pseudogene (red); the analysis included only species for which no morphological data were missing. See Table 1 for the coefficients of each variable in each vector. Vertical and horizontal lines show mean scores for the two species groups **ALT TEXT:** A scatter plot of two sexually selected traits based on results from the clustering analysis.

The evolution of small male size was a good predictor of the evolution of a pseudogene. The phylogenetic logistic regression coefficient for relative male mass was negative (*b* = -4.3e-4), and the standardized effect was large (*z* = -2.17) and significant (*p* = .029). That result was consistent with the one-sample test. Among 11 lineages with a *ZP3R* pseudogene, mean relative male mass (range) was -2.9 kg (−15.6 to 3.1 kg) (i.e., males were smaller than expected) (Figure 4). By contrast, the phylogeny-wide expected relative male mass (range) was 1.3 kg. That difference was in the same direction as the phylogenetic logistic regression and was also significant (*t* = -2.52, *df* = 10, *p* = .03).

**Figure 4.**
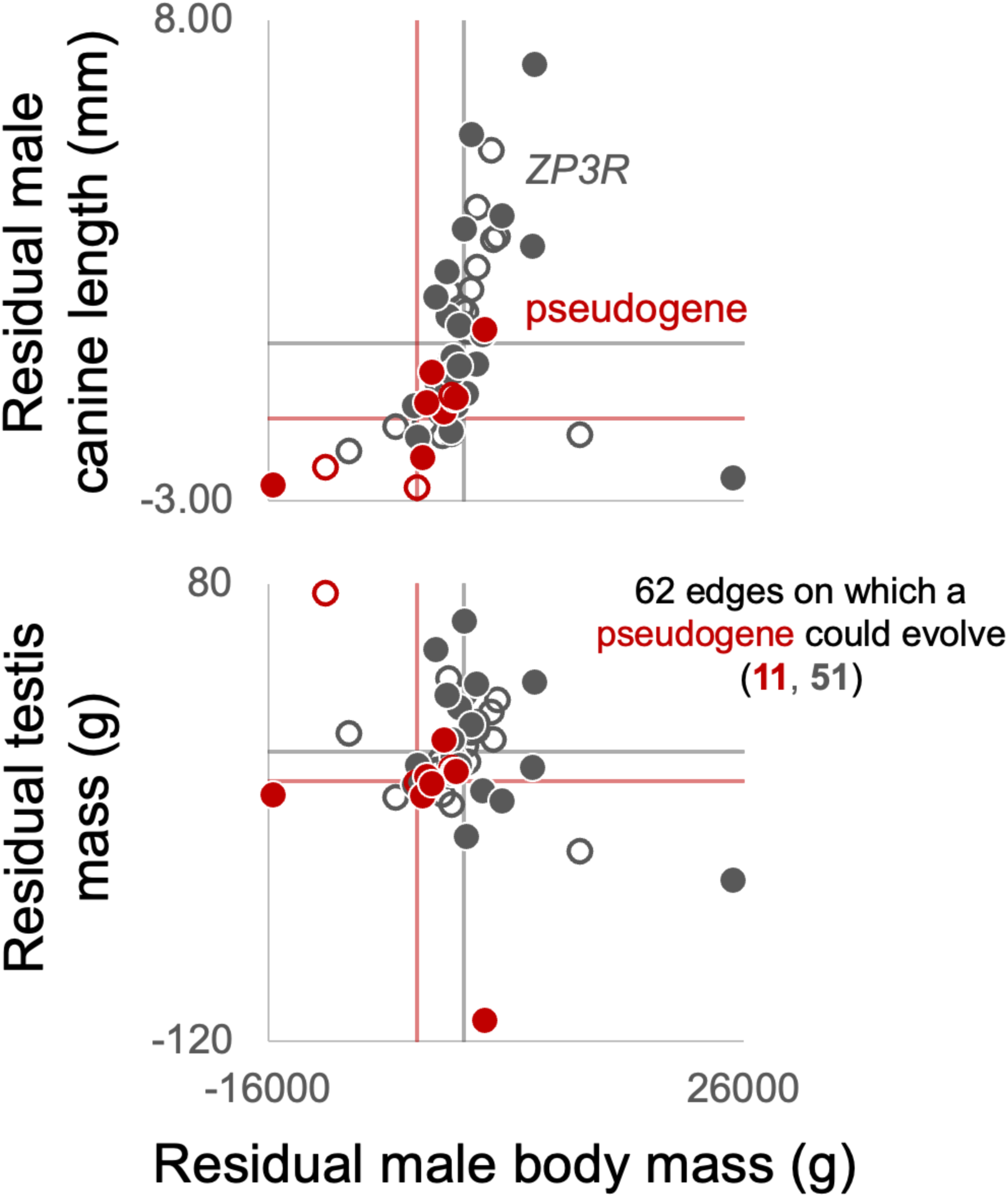
Relative male body mass, relative male canine length, and relative testis mass estimated as residuals from linear regressions. Each symbol shows values for a terminal edge of the phylogeny leading to one of 29 primate species (closed symbols) or values inferred using BayesTraits for one of 33 internal edges of the phylogeny (open symbols). Vertical and horizontal lines show mean values for the 11 total edges on which a pseudogene evolved (red) or the phylogeny-wide expected value on all other edges with a functional *ZP3R* (the known ancestral state). Species and edges included in the analysis are shown in Figure 1 **ALT TEXT:** Two scatter plots of sexually selected traits based on estimates of ancestral trait values from the primate phylogeny.

Among the three variables, small male canines were the strongest predictor of the evolution of a pseudogene. The phylogenetic logistic regression coefficient was negative and the effect of relative male canine length on pseudogenization was significant (*b* = -0.737, *z* = -2.37, *p* = .017). In 11 lineages with a *ZP3R* pseudogene, mean relative male canine length was -1.1 mm (−2.7 to 0.92 mm) (i.e., male canines were smaller than expected) (Figure 4). By contrast, the phylogeny-wide expected canine size was 0.60 mm (Figure 4). In the one-sample test, that large difference in canine size (−1.71 mm) between lineages with a pseudogene and the phylogeny-wide expected value was highly significant (*t* = -4.97, *df* = 10, *p* < .001).

Those two patterns among measures of relative male body mass and relative male canine length were also consistent across the four major primate lineages in our dataset. A *ZP3R* pseudogene evolved 3 or 4 times each among apes, colobines, and New World monkeys with relatively small males and small canines (Figure 5). Those patterns were consistent despite the broad range of male sizes among apes and the range of canine sizes among macaques and baboons. Among the apes, pseudogenes arose mainly in lineages with small males and small canines (humans, chimps, gibbons). No pseudogene evolved among the macaques and baboons, which all have large males with large canines.

**Figure 5.**
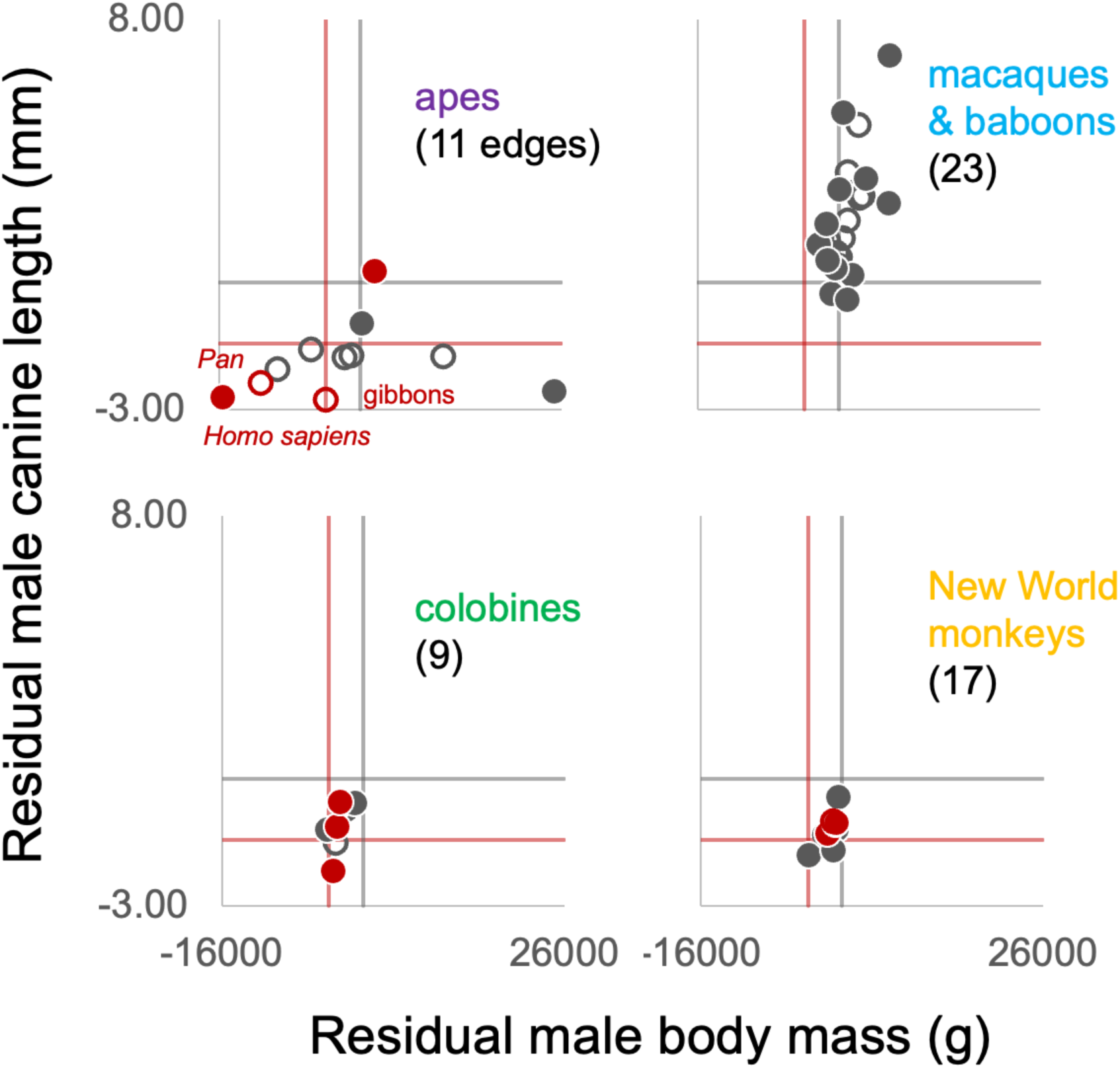
Relative male body mass and relative male canine length from Figure 4 disambiguated by the four major primate lineages (see Figure 1). Each symbol shows values for a terminal edge on the phylogeny leading to one of 29 primate species (closed symbols) or values inferred using BayesTraits for one of 33 internal edges of the phylogeny (open symbols). Vertical and horizontal lines show mean values for the 11 total edges on which a pseudogene evolved (red) or the phylogeny-wide expected value on all other edges with a functional *ZP3R* (the known ancestral state). Species and edges included in the analysis are shown in Figure 1. **ALT TEXT:** For scatter plots of relative male canine length plotted against relative male body mass shown separately for the apes, the macaques and baboons, the colobines, and the New World monkeys.

Surprisingly, testis size did not predict pseudogenization. Mean relative testis mass was similar among lineages that did (−6.22 g) and did not (6.70 g) evolve a pseudogene, relative to the broad scale of relative testis sizes among the species in our sample (range -110 g to 75 g), and there was no obvious relationship between relative testis mass and relative male body mass (Figure 4). The phylogenetic logistic regression coefficient was small, not significantly different from zero (*b* = -8.66e-5, *z* = 0.0162, *p* = .98), and similar to the results from the one-sample test (*t* = -1.01, *df* = 10, *p* = .34).

## Discussion

We found that a *ZP3R* pseudogene tends to evolve in lineages that have evolved smaller values for sexually-selected male traits. A reasonable interpretation of those patterns is that a change in the strength of sexual selection on male competition for access to females (before copulation) or access to eggs (after copulation) allows for the evolution of relatively smaller male body and canine sizes, and that such circumstances also permit or favour the loss of gene function in systems with redundancy in sperm adhesion to the egg coat. We conclude that loss of *ZP3R* function may also be a consequence of relaxed sexual selection on males. The patterns are similar to the insertion of regulatory regions that enhance sex-specific gene expression among some of the same primates in response to changes in sexual selection inferred from analysis of the same three morphological traits by Anderson & Jones (2024).

Carlisle et al. (2024) showed that *ZP3R* evolves under positive selection (a high relative rate of nonsynonymous substitutions) among primate species with a functional gene. They argued that the parallel origins of pseudogenization, the divergence of functional genes under positive selection, and fixation of the human pseudogene mutation all argue against neutral processes (such as relaxed selection in humans and other lineages) and favour some form of strong selection as the cause of pseudogenization. Our results are consistent with that argument and suggest that sexual selection causes both the loss of *ZP3R* function (in species that are experiencing weaker sexual selection or have resolved sexual conflicts) and amino acid sequence divergence among functional *ZP3R* genes (in species that are experiencing strong sexual selection on males or sexual conflict between males and females; Gavrilets et al., 2000).

We found consistent quantitative associations between male body size or canine size and *ZP3R* pseudogenization, but less evidence linking relative testis size and *ZP3R* evolution. Those mixed results suggest that some other traits might account for among-lineage variation in the selective conditions that favour loss of *ZP3R* function in fertilization. One good candidate is behavioural interactions that result in monogamous male-female pairs or that permit single males to monopolize matings with several females (Harcourt et al., 1995; Lindenfors & Tullberg, 1998; Cassini, 2020). Some of our observations are consistent with a role for such traits in accounting for variation among some primate lineages with different mating systems and potential for sperm competition, sexual selection among males, and sexual conflicts of interest between mates. Future studies could also extend the comparison of *ZP3R* structure and life history variation to the lemurs and lorises, in which complex patterns of variation in life history and mating system traits (Kappeler, 1997; Kappeler et al., 2017) are expected to parallel those found among apes, monkeys, and tarsiers and predict the evolution of *ZP3R* structure and function. Unfortunately, we were not able to find and unambiguously annotate *ZP3R* in available genomes from lemurs and lorises (Shao et al., 2023).

We emphasize that these phenotypic traits associated with behavioural interactions between males or between mates may predict *ZP3R* pseudogenization but the causal mechanism of *ZP3R* pseudogenization is more likely to involve the number, durability, and timing of molecular interactions between sperm and eggs at fertilization. These molecular interactions are cryptic in primates but can be observed and quantified in aquatic animals with planktonic gametes and fertilization (Evans & Lymbery, 2020). Changes in the ecology of gamete interactions in the ocean cause the molecular evolution of sperm–egg binding genes (e.g., Levitan, 2012; Patiño et al., 2016). Comparable mechanisms based on the ecology of sperm–egg interactions (e.g., Weber & Fisher, 2023) are likely to cause the evolutionary loss of gene expression such as the repeated parallel pseudogenization of primate *ZP3R*. We know of few previous studies (e.g., Goudet et al., 2008) that have discovered specific genes lost from a fertilization repertoire (as in Carlisle et al., 2024) and then tried to identify features of the selective environment that are associated with loss of gene function and expression (as in this study). But incidental discoveries of loss of gene expression in comparative transcriptomic studies of species with large qualitative differences in life history traits (e.g., species with external versus internal fertilization; Guerra et al., 2021) suggest that loss of function may also be common in comparisons among species with quantitative differences in mating systems (e.g., primates with stronger or weaker precopulatory competition for mates; Carlisle et al., 2022; Rivera & Swanson, 2022).

Carlisle et al. (2024) showed that *ZP3R* originated from a gene duplication of *C4BPA* in a common ancestor that was older than the divergence between Glires and Primates. Genes with other functions can be recruited into gamete binding, possibly as evolutionary solutions to complement or overcome adaptations in the other sex in the context of sexual conflicts of interest (Gavrilets, 2000) over the frequency or intensity of mating or reproductive allocation (Carlisle et al., 2022; Rivera & Swanson, 2022). How old was this gene duplication event? Extending these analyses to the other major lineages of mammals could identify the age and origin of *ZP3R* and increased redundancy in gene function at fertilization.

These future studies could also help to understand the causes of decreased redundancy in gene function when *ZP3R* (and other fertilization genes) is lost by pseudogenization. Our analysis suggests that altered sexual selection is associated with this phenomenon. That proposed relationship implies an evolutionary cost to maintaining redundancy in the suite of primate fertilization genes under conditions of relaxed sexual selection on males or resolution of sexual conflict between mates. The nature of this cost is not known but might include high risk of fatal polyspermy relative to the reduced benefits of high-fidelity sperm binding in species with lower sperm competition among males (Okamoto, 2016; Levitan et al., 2019; Weber & Fisher, 2023; but see Dale, 2024). Under this hypothesis, males in species that have resolved sexual conflict or reduced male competition under less promiscuous mating systems may experience selection for lower sperm binding affinity to the egg coat and reduced likelihood that two sperm from the same male fertilize the same egg (resulting in the deaths of all three genome copies).

Removing some male-expressed loci from the egg-binding repertoire of sperm could be one response to such selection, similar to changes in allele frequencies for single loci in response to changes in the male optimum for rates of sperm–egg contact (Levitan, 2012). That hypothesis also predicts that, in species that have not evolved this kind of resolution of sexual conflict, females may also experience selection for lowering sperm binding affinity to the egg coat by adding new female-expressed genes to the egg coat repertoire in order to fend off supernumerary sperm and avoid polyspermy (e.g., Aagaard et al., 2013).

Reducing the size of testes and the number of sperm per ejaculate seems like another obvious potential response to selection on males to avoid polyspermy, and might be associated with genomic changes that reduce gene expression that mediates sperm– egg binding. Anderson & Jones (2024) also found phylogenetic correlations between changes in testis size and the number of genomic elements that affect androgen-dependent gene expression. By contrast, our results do not suggest that adaptive change in testis size is often associated with the evolution of a *ZP3R* pseudogene, and we are not able to explain why the evolution of some sexually selected male traits (body size, canine length) but not others (testis size) might be associated with this recurrent gene loss. Lupold et al. (2019) reviewed the evidence for a tradeoff between male investment in weapons, badges of status, or other mechanisms of precopulatory mate competition versus investment in testis size for postcopulatory sperm competition. One speculative hypothesis is that testis size (and number of sperm) may show a similar evolutionary tradeoff with molecular traits of sperm (and the expression of genes that moderate the affinity of each sperm for the egg coat).

The causes of decreased redundancy in human fertilization (through the evolution of the human *ZP3R* pseudogene) are of particular interest. The timing of the human pseudogenization event could point to its evolutionary cause. Because the premature stop codon in the second CCP domain of human *ZP3R* is not a polymorphism in dbSNP (Sherry et al., 1999), and because both the divergence and loss of *ZP3R* among primates appears to be driven by selection (Carlisle et al., 2024), the origin and fixation of the human pseudogene is expected to be associated with a change in selection acting on the early human lineage. Among the Neanderthal and Denisovan genomes in the UCSC genome browser (GRCh37/hg19), three Neanderthal sequence reads and 25 Denisovan reads map to the same site (chr1:207340753) as the human stop codon in *ZP3R*; all have the human TAA stop mutation. That similarity relative to the depth of coverage suggests that the human pseudogene is shared between archaic and early modern humans, and that the causes of the human pseudogenization event were older than the divergence between them and unrelated to any life history or mating system differences among them.

## Author contributions

JAC compiled and analyzed data, designed the figures, and wrote the paper; ALB compiled and analyzed data and wrote the paper; MWH conceived the project, compiled and analyzed data, designed the figures, and wrote the paper.

## Acknowledgments

Members of the FAB/Crawford lab at Simon Fraser University provided constructive criticism of the project. David Fisher, Carlos Schrager, Jon Slate, and anonymous reviewers provided helpful reviews that greatly improved on early versions of the study and manuscript. Thanks to Doug Brandon, Mark Collard, Maria Creighton, Mick Elliot, Dan Greenberg, Michael Krützen, Ruth Linsky, Fanny Mazzamurro, Brea McCauley, Mark Skinner, Larry Ulibarri, and Rutger Vos for help with data and analyses.

## Funding

The data analysis and writing were supported by Natural Sciences and Engineering Research Council Discovery Grant R611839 to MWH.

## Conflict of interest statement

The authors declare no conflicts of interest.

## Data and code availability

All data analyzed in this study are available as supplementary data files.

## Supplementary Data

Appendix S1. Supplementary methods for identifying *ZP3R* pseudogenes in 17 genomes from Shao et al. (2023).

Appendix S2. Data and sources for five morphological variables in 48 primate species (including incomplete data for 12 species).

Appendix S3. Complete data for five morphological variables in 36 primate species, with R code and results for the principal components analysis (eigenvalues, eigenvectors, scores for each species).

Appendix S4. Phylogenetic logistic regression analyses of *ZP3R* pseudogenization as a function of relative male size.

Appendix S5. BayesTraits inferences of relative male size on internal edges of the primate phylogeny.

## Notes

### Competing Interest Statement

The authors have declared no competing interest.

### Summary of Updates

We added new data and analyses, including new ZP3R gene copies identified from other primate genomes. To do that bioinformatics work we recruited a new collaborator and coauthor (Dr. Carlisle). We used improved comparative methods (phylogenetic logistic regressions) to analyze the association between pseudogenization and life history traits.

