## Appendix S1 for "Sexual selection and pseudogenization in primate fertilization": Supplementary_methods.pdf

### Supplementary methods: Identification of *ZP3R* and *C4BPA* in genomes from Shao et al. (2023) and determination of pseudogenization status

Shao et al. (2023) published new reference genomes for 27 primate species. We used the following workflow to identify additional primate *ZP3R* sequences for analysis along with the genomes investigated in Carlisle et al. (2024). The updated results are illustrated in the image file called "Shao_Primates_ZP3R_FigureS1.png" (next page).

1. Query the Shao et al. (2023) genomes using orangutan (*Pongo abelli*) amino acid sequence for *ZP3R* and *C4BPA* using tblastn

Using the orangutan amino acid sequences from Carlisle et al. (2024), we used tblastn to identify scaffolds in each new primate species genome that may contain either or both genes. In all species investigated in Carlisle et al. (2024) (rodents and primates), these genes exist as tandem paralogs. These proteins are close paralogs, and can be difficult to distinguish from each other, so we collected sequences for both. We used orangutan *ZP3R* because it is the only ape we investigated in Carlisle et al. (2024) that has a non-pseudogenized *ZP3R*. We also used tblastn to identify genes that are found upstream or downstream of *ZP3R* and *C4BPA* in orangutans in order to establish the syntenic context for those two genes relative to genes expected to be found in the same region of chr1 (*YOD1*, *C4BPB*, *PFKFB2*, *CD55*).

The result of these tblastn queries are recorded in the supplementary file "Shao_Collection.xlsx" with a tab for each species.

In some cases, the top hit for *C4BPA* (and a couple of times *ZP3R*) was on a different scaffold than its paralog and its syntenic genes, and the second-best hit was on the scaffold with the top hit for *ZP3R* and syntenic genes. We collected the scaffolds for the top 2 tblastn hits (for hits that exist within the syntenic locus or not) for *C4BPA* and *ZP3R* for follow-up analysis.

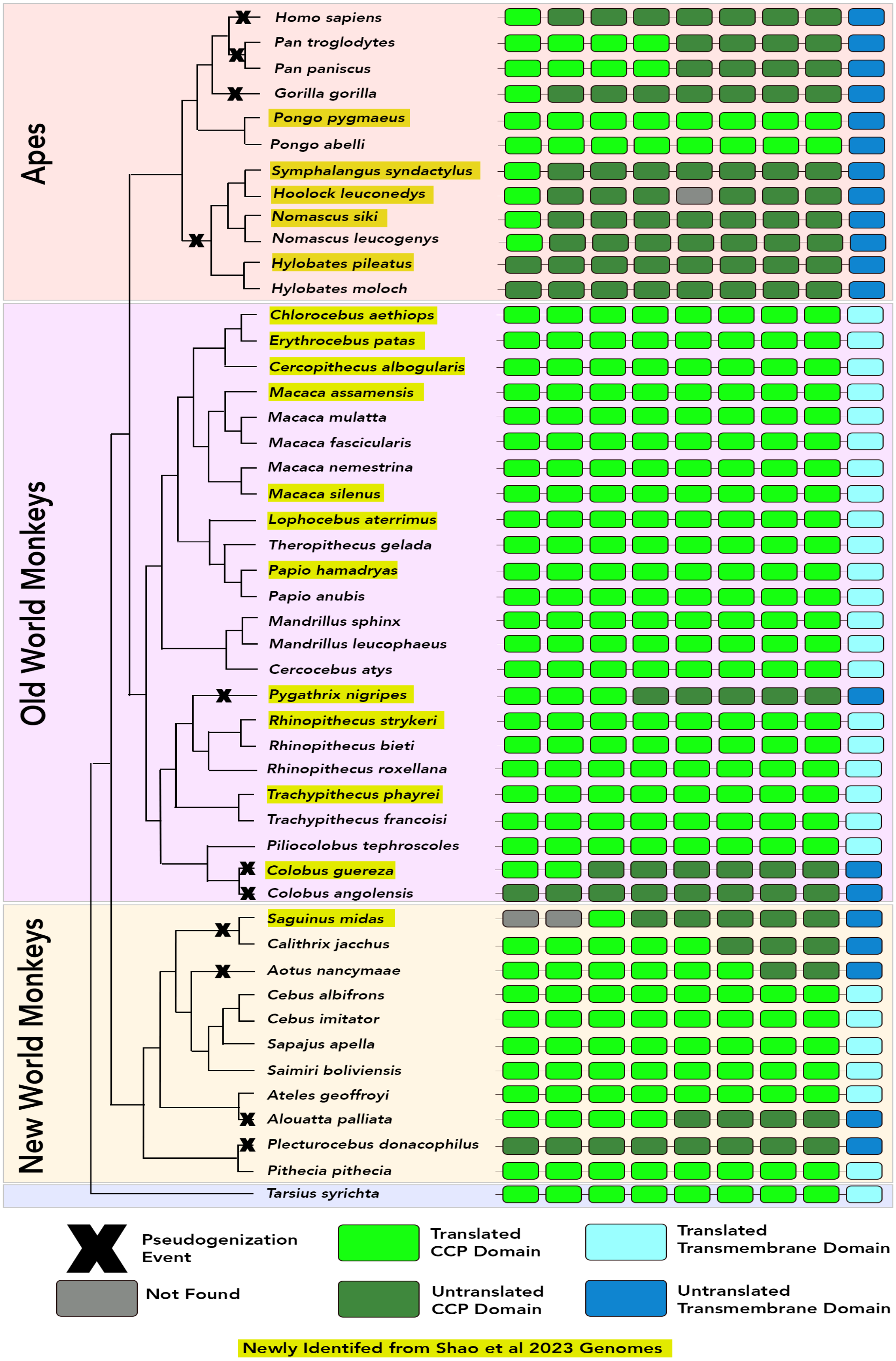
Apes
Old World Monkeys
New World Monkeys
Homo sapiens
Pan troglodytes
Pan paniscus
Gorilla gorilla
Pongo pygmaeus
Pongo abelli
Symphalangus syndactylus
Hoolock leuconedys
Nomascus siki
Nomascus leucogenys
Hylobates pileatus
Hylobates moloch
Chlorocebus aethiops
Erythrocebus patas
Cercopithecus albogularis
Macaca assamensis
Macaca mulatta
Macaca fascicularis
Macaca nemestrina
Macaca silenus
Lophocebus aterrimus
Theropithecus gelada
Papio hamadryas
Papio anubis
Mandrillus sphinx
Mandrillus leucophaeus
Cercocebus atys
Pygathrix nigripes
Rhinopithecus strykeri
Rhinopithecus bieti
Rhinopithecus roxellana
Trachypithecus phayrei
Trachypithecus francoisi
Piliocolobus tephroscoles
Colobus guereza
Colobus angolensis
Saguinus midas
Calithrix jacchus
Aotus nancymaae
Cebus albifrons
Cebus imitator
Sapajus apella
Saimiri boliviensis
Ateles geoffroyi
Alouatta palliata
Plecturocebus donacophilus
Pithecia pithecia
Tarsius syrichta
Pseudogenization Event
Not Found
Translated CCP Domain
Untranslated CCP Domain
Translated Transmembrane Domain
Untranslated Transmembrane Domain
Newly Identifed from Shao et al 2023 Genomes

2. Use Exonerate to identify predicted protein sequences for C4BPA and ZP3R for each species.

(https://www.ebi.ac.uk/about/vertebrate-genomics/software/exonerate-manual)

Using the orangutan amino acid sequences as queries, we used the protein2genome function of Exonerate to extract the top predicted CDS of each of these proteins from the scaffolds identified in the prior step.

We translated the top predicted CDSs to amino acid sequences. The top scoring Exonerate predicted CDS sequence for each gene was in all cases from the scaffold identified in step 1 that also contained the syntenic genes (see "Shao_Collection.xlsx"). This was true even if the region identified by tblastn in step 1 that was non-syntenic had a better E-value; we think this may be caused by exon-intron structure affecting the significance of tblastn hits in step 1. The results for each species are collected in the directory called "Exonerate".

3. Use tblastn to search putative ZP3R and C4BPA protein sequences against the *Pongo abelli* transcriptome.

We queried the sequences extracted in step 2 against the transcriptome of *Pongo abelli* using tblastn (reciprocal blast). For the protein sequences found in the non-syntenic region, the top hit was *SVEP1* in orangutans. For the syntenic genes, the top hits were either *ZP3R* or *C4BPA*. Despite not always being the top hit from the initial tblastn search in step 1, these results indicate that the syntenic sequence is more likely to be the true *ZP3R* or *C4BPA* sequence.

We also used reciprocal best blast to confirm the identity of the genes surrounding *ZP3R* in the *Pongo abelli* genome (*YOD1*, *C4BPB*, *PFKFB2*, *CD55*) to establish a syntenic location for *ZP3R*. These results are collected in the supplementary file called "All_Annotated_Scaffolds.gff".

4. Clustal Omega alignments of newly identified *ZP3R* and *C4BPA* sequences to compare sequence identity to orangutan (*Pongo abelli*) *ZP3R* and *C4BPA*.

Using Clustal Omega, we compared the sequence identity of the putative *ZP3R* and *C4BPA* hits to orangutan sequences. The syntenic hits showed a higher sequence identity to either *ZP3R* or *C4BPA*. This supports the identification of the syntenic sequences as the orthologs of *ZP3R* and *C4BPA* (see "Shao_Collection.xlsx"). On the basis of all of those results (synteny, Exonerate prediction score, tblastn comparison to the *Pongo abelli* transcriptome, and sequence identity), we eliminated the sequences from the non-syntenic genomic region as potential *ZP3R* or *C4BPA* orthologs. Instead, we tentatively identify those other hits as probable *SVEP1* orthologs.

5. Phylogenetic clustering of collected C4BPA and ZP3r amino acid sequences to resolve orthology vs paralogy

We created a protein alignment that included all the C4BPA and ZP3R sequences from rodents and primates analyzed in Carlisle et al. (2024) plus the newly identified candidate C4BPA and ZP3R protein sequences from our analyses of the genomes in Shao et al. (2023). We used raxml-ng to construct a phylogeny (1000 bootstraps). This is the same approach used to create Figure 1B in Carlisle et al. (2024) to resolve paralogy vs orthology (the same raxml-ng parameters were also used as in Carlisle et al 2024 for protein phylogeny construction). The results are summarized in "Shao_Collection.xlsx".

For 5 species, we were not able to clearly identify ZP3r (potentially for assembly issues). These species failed at various steps of our pipeline. Those species were not included in our subsequent analyses described in the main text.

6. Identification of unique pseudogenization events via parsimony

The Exonerate output files for identify pseudogenizing mutations (premature stop codons, frameshift insertions or deletions). For all new *ZP3R* sequences with a pseudogenizing mutation, we used MAFFT in Geneious to create a protein alignment (pseudogenizing mutations corrected) and annotated the predicted pseudogenizing mutations onto the alignment with respect to the locations of the predicted CCP domains. Using this alignment, we were able to visualize shared vs unique pseudogenizing mutations. Based on parsimony, we were able to determine whether closely related primates (e.g., the six gibbons, or the two species of *Colobus*) with a pseudogenized *ZP3R* were likely to have inherited the pseudogene from their common ancestor (one origin) or more likely to have evolved a pseudogene in parallel (two or more origins). The sequences and alignments are included in the directory called "Sequence Files". The genomic locations of the observed pseudogenizing mutations in all species with a pseudogene are summarized in the file called "Alignment of Pseudogenized ZP3R Protein Sequences in Primates.png" (below).

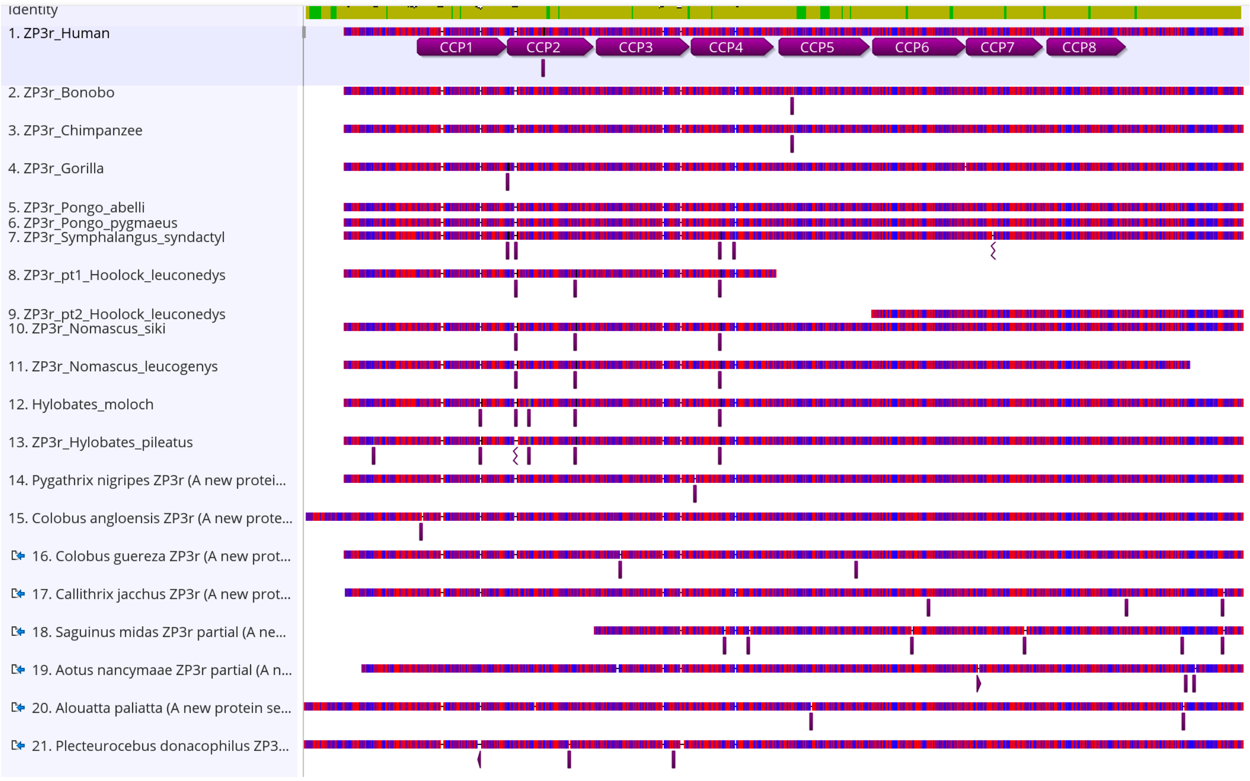
Identity
1. ZP3r_Human
CCP1
CCP2
CCP3
CCP4
CCP5
CCP6
CCP7
CCP8
2. ZP3r_Bonobo
3. ZP3r_Chimpanzee
4. ZP3r_Gorilla
5. ZP3r_Pongo_abelli
6. ZP3r_Pongo_pygmaeus
7. ZP3r_Symphalangus_syndactyl
8. ZP3r_pt1_Hoolock_leuconedys
9. ZP3r_pt2_Hoolock_leuconedys
10. ZP3r_Nomascus_siki
11. ZP3r_Nomascus_leucogenys
12. Hylobates_moloch
13. ZP3r_Hylobates_pileatus
14. Pygathrix nigripes ZP3r (A new protei...
15. Colobus angloensis ZP3r (A new prote...
16. Colobus guereza ZP3r (A new prot...
17. Callithrix jacchus ZP3r (A new prot...
18. Saguinus midas ZP3r partial (A ne...
19. Aotus nancymaae ZP3r partial (A n...
20. Alouatta paliatta (A new protein se...
21. Plecteurocebus donacophilus ZP3...
