## Supplementary figures and images for "Sexual selection and pseudogenization in primate fertilization"

### Alignment of Pseudogenized ZP3R Protein Sequences in Primates.png

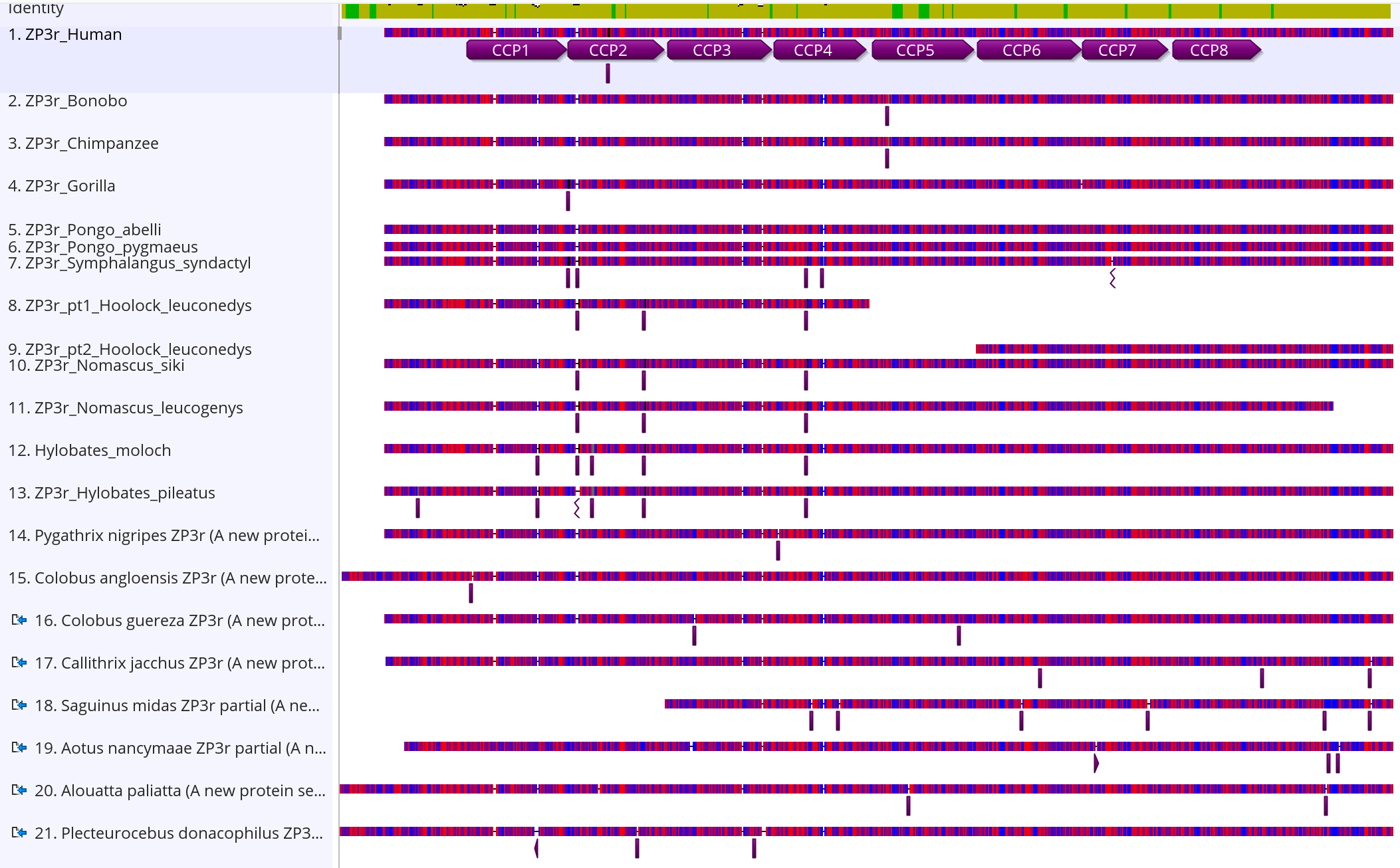

### Shao_Primates_ZP3r_Figure1.png

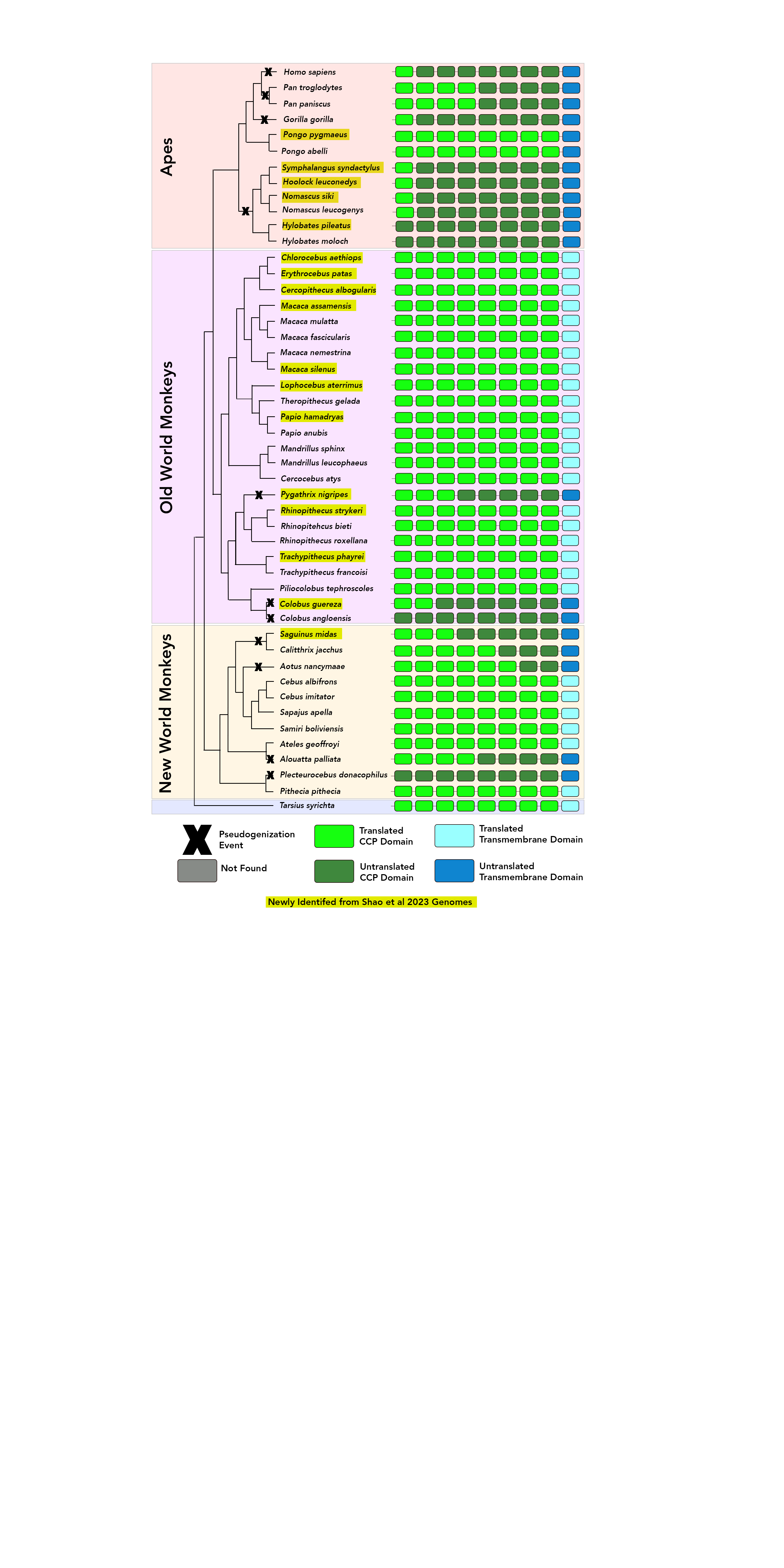

### Shao_Primates_ZP3r_Figure1_v2_missing.png

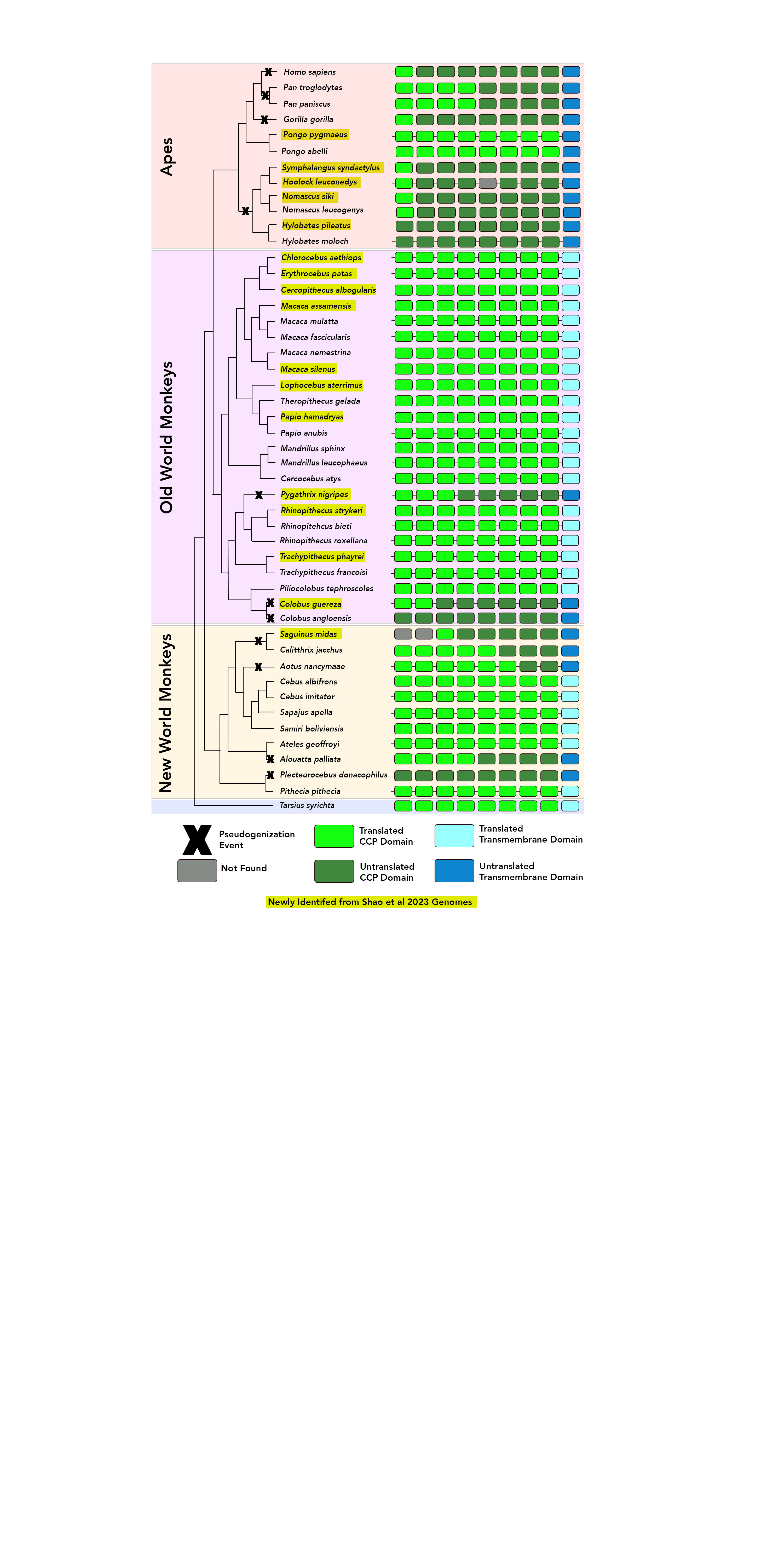

### Shao_Primates_ZP3R_FigureS1.png

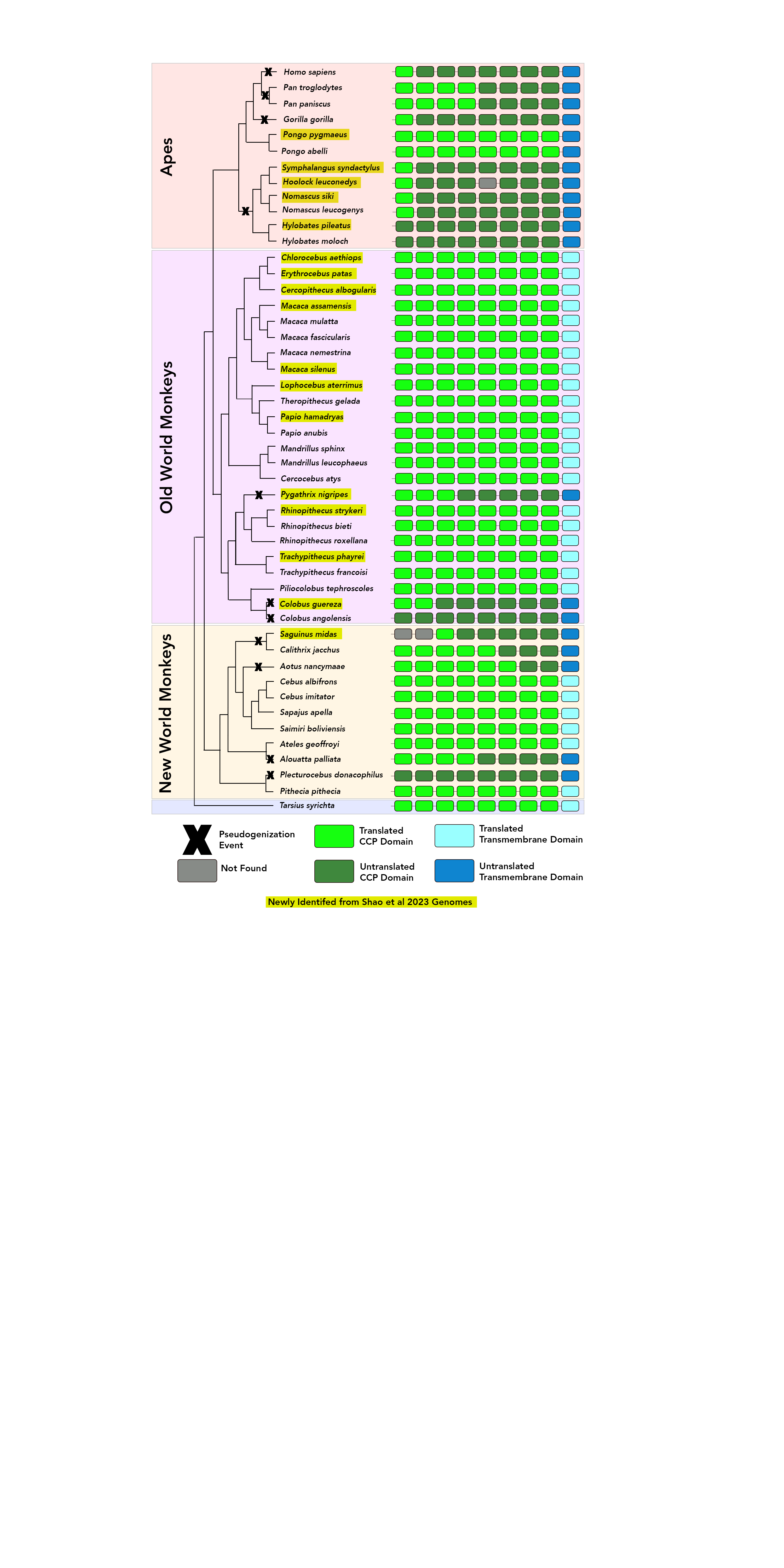
